# EEG Alpha Power Predicts the Temporal Sensitivity of Multisensory Integration

**DOI:** 10.1101/2020.08.26.268144

**Authors:** Raquel E. London, Christopher S. Y. Benwell, Roberto Cecere, Michel Quak, Gregor Thut, Durk Talsma

## Abstract

Pre-stimulus EEG oscillations, especially in the alpha range (8-13 Hz), can affect the integration of stimulus features into a coherent percept. The effects of alpha power are often explained in terms of alpha’s inhibitory functions, whereas effects of alpha frequency have bolstered theories of discrete perceptual cycles, where the length of a cycle, or window of integration, is determined by alpha frequency. Such studies typically employ visual detection paradigms with near-threshold or even illusory stimuli. It is unclear whether such results generalize to above-threshold stimuli. Here, we recorded electroencephalography, while measuring temporal discrimination sensitivity in a temporal order judgement task using above-threshold auditory and visual stimuli. We tested whether pre-stimulus oscillations predict audio-visual temporal discrimination sensitivity on a trial-by-trial basis. By applying a jackknife procedure to link single-trial pre-stimulus oscillatory power and instantaneous frequency to psychometric measures, we identified two highly overlapping clusters over posterior sites. One where lower alpha power was associated with higher temporal sensitivity of audiovisual discrimination, and another where higher instantaneous alpha-frequency predicted higher temporal discrimination sensitivity. A follow-up analysis revealed that these effects were not independent, and that the effect of instantaneous frequency could be explained by power modulations in the lower alpha band. These results suggest that temporal sensitivity for above-threshold multisensory stimuli changes spontaneously from moment to moment and is likely related to fluctuations in cortical excitability. Moreover, our results caution against interpreting instantaneous frequency effects as independent from power effects if these effects overlap in time and space.

## Introduction

A fundamental aspect of perception is the integration of sensory signals to form meaningful percepts within and across modalities. Whether two signals from different sensory modalities are integrated depends, among other factors, on their temporal proximity. The shorter the time between them, the higher the chance they will be integrated (Lewald & Guski, 2003; Meredith et al., 1987; Senkowski et al., 2007).

Exactly how close in time these signals need to be for integration to occur depends on several factors. Accordingly, temporal sensitivity is highly variable both within and between individuals. For example, in people diagnosed with schizophrenia, autism or dyslexia, audiovisual temporal sensitivity appears to be reduced compared to healthy controls (De Boer-Schellekens et al., 2013; Foucher et al., 2007; Hairston et al., 2005; Martin et al., 2013; Stevenson et al., 2012, 2014, 2017; Wallace & Stevenson, 2014). Even in the healthy population, the temporal sensitivity of integration differs markedly across individuals (Stevenson et al., 2012). Within individuals, temporal sensitivity is modulated by factors such as stimulus complexity (Stevenson & Wallace, 2013), stimulus intensity (Fister et al., 2016) and spatial relation (Lewald & Guski, 2003). Previous experience and training (Lee & Noppeney, 2011; Navarra et al., 2005; Powers et al., 2009), attention (Donohue et al., 2015; Talsma et al., 2009) and cognitive load (Dean et al., 2017) also have an impact.

In the search for the neural correlates of temporal sensitivity in multisensory integration, most previous studies, including some of those discussed above, have focused on transient, stimulus-related activity. Recently, however, it has been found that spontaneous oscillatory EEG activity reflecting momentary state can affect temporal discrimination sensitivity in unisensory auditory, tactile and visual perception (Baumgarten et al., 2016; Bernasconi et al., 2011; Samaha & Postle, 2015). In the multisensory domain, the relationship between ongoing oscillatory brain activity and multisensory integration has increasingly been studied as well (Cecere et al., 2015; Grabot et al., 2017; Ikumi et al., 2019; Keil & Senkowski, 2017; Leonardelli et al., 2015; Ronconi et al., 2018; Yuan et al., 2016).

For example, Cecere et al. (2015) used the sound-induced flash illusion in combination with EEG and transcranial alternating current stimulation to show that peak alpha-frequency around stimulus presentation causally determined the temporal window of audiovisual integration. Their findings were corroborated by (Keil & Senkowski, 2017) based on an analysis of pre-stimulus alpha activity alone. These results are informative regarding the temporal characteristics of auditory influences on visual perception (sound-induced flash illusion), but the question remains whether they are representative of general multisensory perceptual processes. Not all participants report the associated illusion and whether they do so might depend on the power of their alpha oscillations (Cecere et al., 2015; Lange et al., 2013); however see Keil & Senkowski, 2017). Furthermore, the effect of alpha oscillations on the temporal sensitivity of perception may be so subtle that it only becomes apparent when the stimuli are around threshold or that it even relies on illusory perception (see Benwell et al., 2017; Iemi & Busch, 2018, for a link between pre-stimulus alpha activity and subjective rather than objective measures of perception).

To test more directly whether spontaneous pre-stimulus activity affects audiovisual temporal sensitivity, we asked participants to make temporal order judgements on suprathreshold audio-visual stimuli. We employed a “jackknife” procedure adapted for linking psychophysical data to single-trial EEG parameters (Benwell et al., 2018; Gluth & Meiran, 2019). This leave-one-out procedure allowed us to examine cross-trial co-variation of pre-stimulus oscillatory parameters in EEG with temporal discrimination sensitivity estimates obtained from psychometric curves. By these means, we tested whether the power of pre-stimulus oscillations (2 to 45 Hz) was predictive of the temporal sensitivity of multisensory integration. Furthermore, to extend the results of Cecere et al. (2015) and (Keil & Senkowski, 2017), we tested whether the instantaneous frequency of pre-stimulus oscillations in the alpha range was positively correlated with individuals’ audio-visual temporal sensitivity.

## Method

### Participants

Forty-three volunteers participated in this experiment for monetary compensation. Two participants were excluded due to their estimated sensitivity measure exceeding the maximum SOA of 350 ms. One participant was excluded due to not completing the experiment. Analyses were carried out on the data of the remaining 40 participants (30 female, 2 left handed, median age: 23, age range: 18 – 32). Participants reported having normal audition and normal or corrected-to-normal vision and no history of neurological disorder or recent use of psychoactive substances. The experiment was approved by the Ethics Committee of Ghent University. Participants gave informed consent prior to the start of the experiment and were financially compensated for their time.

### Apparatus and Stimuli

Participants were seated in a dimly lit, sound-proof and electrically shielded chamber, with their head stabilized by a chinrest at 50 cm from a 24-inch LCD monitor (BenQ XL2411; 120 Hz refresh rate). The task was an audiovisual temporal order judgement task (TOJ) in which participants were presented with a visual flash and an auditory beep, and then asked to judge which of the two had been presented first (See figure 1). The experiment had one within-participants factor which was the stimulus onset asynchrony (SOA) between the flash and the beep. The SOA had 12 levels (−350, −216, −133, −88, −50, −16, +16, +50, +88, +133, +216 and +350 ms) where negative SOAs indicate that the beep was presented first and positive SOAs that the flash was presented first. Each SOA was presented 70 times giving a total of 840 trials, divided over 35 blocks of 24 trials each. SOA was randomized per block with each condition presented twice within each block. The task was implemented using the E-prime 1.2 software package (Schneider et al., 2002) on an HP Compaq desktop computer running Microsoft Windows XP. This setup allowed for the timing of stimulus presentation to be at a resolution of <= 1 ms which was confirmed with an oscilloscope. The visual stimulus was a solid white circle (luminance of 270 cd/m^2^) subtending a visual angle of 1.95°. It was presented at 5° below a central fixation cross subtending 0.46° on a black background and for a duration of 16.6 ms. The auditory stimulus was a 1850 Hz tone presented at 72 dB(A) with a duration of 16 ms (plus 3 ms fade-in and 3 ms fade-out) delivered by two loudspeakers (Logic3 Screenbeat ES20). The loudspeakers were placed to the left and right of the monitor.

**Figure 1.**
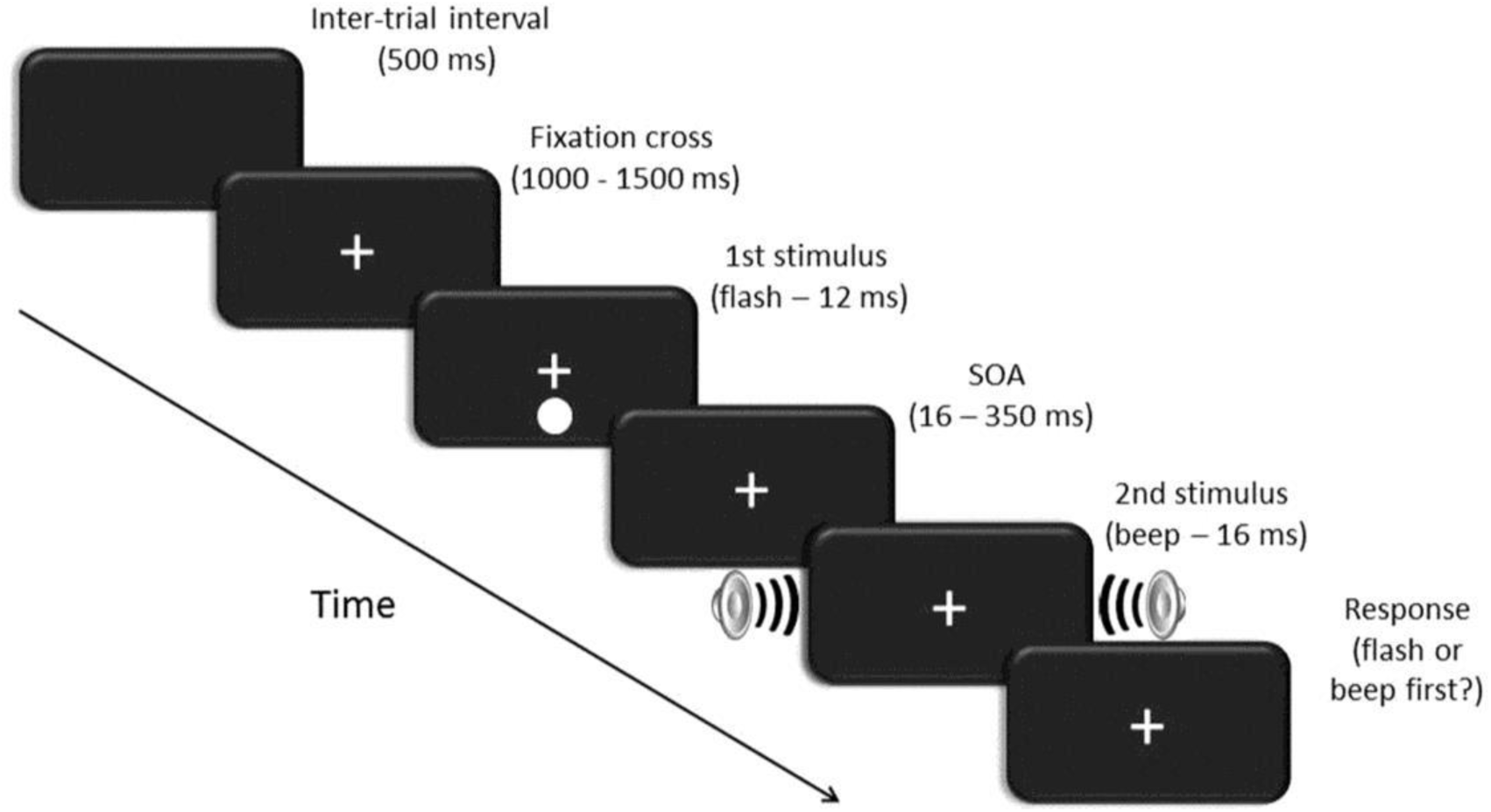
Schematic representation of one experimental trial. After an inter-trial interval of 500 ms, a fixation cross appeared on the screen for a random duration between 1000 ms and 1500 ms. Then the first stimulus (in this case a flash) was presented, followed by the second stimulus (in this case a beep) at one of 12 possible delays (SOAs). The fixation cross remained on the screen until participants had responded which of the two events they had perceived first.

**Figure 2.**
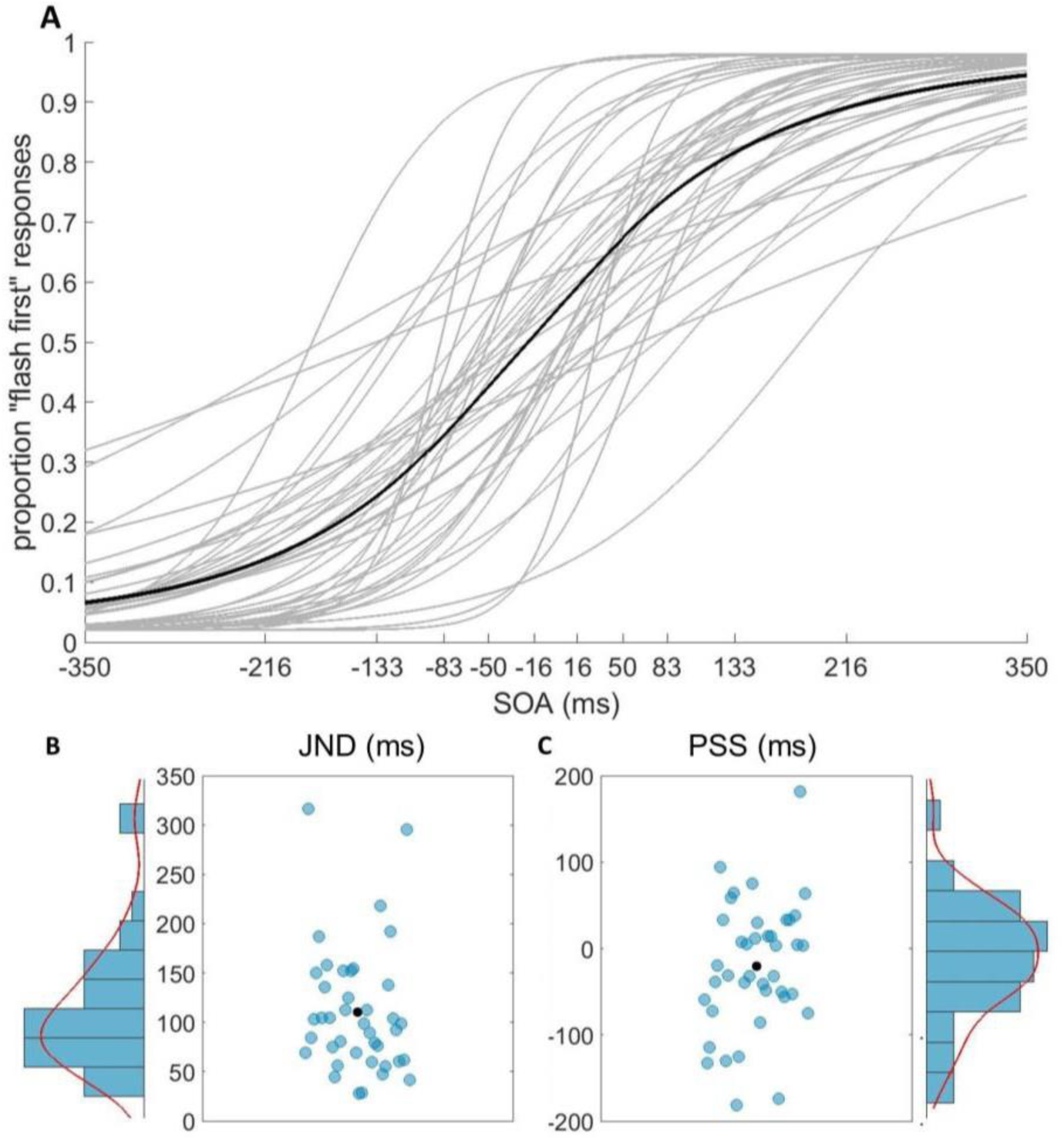
Behavioural results. **(A)** Psychometric functions fit to each participant’s data are displayed in grey (n=40). The psychometric function fit to the group averaged data is displayed in black. **(B)** The JND’s derived from the psychometric function of each participant are displayed in blue. The average JND is displayed in black. **(C)** The PSS’s derived from the psychometric function of each participant are displayed in blue. The average PSS is displayed in black.

### Procedure

The experiment started with the recording of 5 minutes of eyes-open resting state EEG and a seven-minute passive observation task with the sequential presentation of 50 instances of the visual stimulus and 50 instances of the auditory stimulus. Since the EEG data collected during this session are beyond the scope of the current project, they will not be reported here. The TOJ task then started after two practice blocks of 12 trials (one for each SOA, order randomized) during which the experimenter was present to ensure participants understood the instructions. Each trial started with the presentation of a central fixation cross. Participants were instructed to fixate this cross throughout the task. After a random interval of 1000-1500 ms, the first stimulus (a flash or a beep, depending on the condition) was presented. After a random delay, chosen amongst the 12 possible SOAs, the second stimulus was presented. Participants were instructed to judge which stimulus (auditory or visual) had been presented first. The task was not speeded, and there was no time limit, but participants were instructed not to think about the answer for too long. Participants pressed the “z” key when they had perceived the auditory stimulus first, and the “m” key when they had perceived the visual stimulus first with the middle finger of their left and right hand, respectively. After the response, a black screen was presented for 500 ms after which the next trial started. In the practice session, participants received feedback after each trial. During the experimental session no single-trial feedback was given, but after each block the mean accuracy for that block was presented. Between blocks, there was a self-paced break during which participants were encouraged to rest for a short moment. The total duration of the experiment was approximately 50 minutes.

### Electrophysiological recording and pre-processing

The electroencephalogram (EEG) was recorded at 1024 Hz with a Biosemi ActiveTwo system (Biosemi, Amsterdam, Netherlands) with 64 Ag–AgCl scalp electrodes positioned according to the standard international 10–20 system. Additional electrodes were positioned at the outer canthi of both eyes and directly above and below the right eye to acquire horizontal and vertical electro-oculograms (EOG), respectively. Preprocessing was done with custom scripts incorporating functions from the EEGLAB toolbox (Delorme & Makeig, 2004). Data was high-pass filtered using a Hamming windowed sinc FIR filter with the lower edge of the pass band at 0.5 Hz and a cutoff-frequency of 0.25 Hz. Data was low-pass filtered using a Hamming windowed sinc FIR filter with the upper edge of the pass band at 45 Hz and a cutoff-frequency of 50.6 Hz. In preparation for independent component analysis (ICA), data was then cut into 2-second epochs starting 1500 ms before and ending 500 ms after the first stimulus. The epoch mean was subtracted and trials containing unique or very large artefacts were manually discarded. Electrodes exhibiting excessive noise were removed and interpolated. In six participants, this was the case for one electrode and in two participants for two electrodes. Data was then re-referenced to the average of all electrodes (excluding external electrodes) and ICA was run with the EEGLAB “runica” function. Subsequently, the filtered continuous data was re-epoched in preparation for time-frequency analysis to 4-second-long epochs starting 2500 ms before until 1500 ms after the first stimulus onset (exceeding the −1500 to +500ms window of interest to avoid filter artifacts at the edges). As before, the epoch mean was subtracted, bad electrodes were eliminated and the data was re-referenced to the average of all electrodes except the electro-oculogram and mastoid electrodes. Then, the previously obtained ICA weights were applied to this dataset and components reflecting eye movements, blinks or muscle artefacts were projected out of the data. The number of components that was removed per participant ranged from 1 to 10 with a median of 3. Next, eliminated electrodes were interpolated and trials containing artefacts were manually discarded. The percentage of trials that was discarded per participant ranged from 2% to 39%, with a median of 9%. Finally, in order to improve topographic localization, a Laplacian transform was applied through the use of Matlab script accompanying (Cohen, 2014).

### Behavioural analysis

We were interested in the minimum amount of time between the auditory and visual stimuli that was needed in order for each participant to be able to correctly judge the order in which the stimuli had been presented in 75% of the trials. This psychophysical measure of temporal sensitivity is referred to as the “just noticeable difference” (JND) and was derived using the Palamedes toolbox for Matlab (Prins & Kingdom, 2018). First, a logistic function was fit to the proportion of “flash-first” responses as a function of SOA. Guess rate and lapse rate were fixed at 0.02 for each participant. The logistic function is given as:

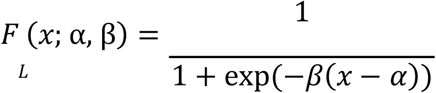

with *x* denoting the SOA, *α* the value of *x* at which the function evaluates to 0.5 and βthe slope or steepness of the function. Second, the difference along the x-axis in milliseconds between 25% and 75% “flash first” responses on the y-axis was divided by two to obtain the JND in milliseconds:

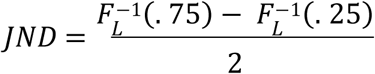

Lower JND values indicate higher temporal sensitivity, whereas higher JND values denote lower sensitivity. Since observations from multiple trials are required to fit a function and derive the JND, this metric is not definable on a single-trial basis. However, it was precisely our aim to investigate how moment-to-moment fluctuations in EEG parameters and temporal sensitivity co-varied across trials. We addressed this challenge by applying a method developed by (Benwell, et al., 2018; see also Gluth & Meiran, 2019) which adapts a “jackknife” procedure (Quenouille, 1949; Richter et al., 2015; Stahl & Gibbons, 2004; Tukey, 1958) to link single-trial variability in oscillatory activity to psychometric measures such as the JND (see an extended explanation of the jackknife analysis below). Another measure that can be derived from the logistic function is the point of subjective simultaneity (PSS). Although this measure is not pertinent to our research question and was not further analyzed in relation to the EEG, we derived it for each participant to offer a more complete description of the behavioural data. The PSS corresponds to the value of the SOA for which the function evaluates to 0.5 and is interpreted as the SOA where the stimuli appear to the observer as arriving simultaneously. The PSS is given as:

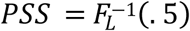

### Electrophysiological analyses: Trial-by-trial variations within participants

#### EEG Power

A time-frequency representation of the single-trial data was obtained by convolving the preprocessed data with a complex wavelet using the “mtmconvol” option of the “ft_freqanalysis” function from the Fieldtrip toolbox (Oostenveld et al., 2011). A sliding window with a length of 500 ms was employed to segment the data. The window was shifted forward in steps of 20 ms. Each segment was multiplied with a Hanning taper to avoid edge artefacts. The value of oscillatory power at each data point therefore included activity from 250 ms before and 250 ms after that time point. Since we were expressly interested in ongoing, stimulus-unrelated activity, we restricted our power analysis to the time points ranging from 750 ms to 250 ms before the onset of the first stimulus to ensure that no stimulus related activity was included. Single-trial oscillatory power was thus obtained at 25 time points in 20 ms steps and 21 frequencies ranging from 2 to 45 Hz in 2 Hz steps for all 64 electrodes. Finally, the resulting power values were converted to decibels.

#### Instantaneous alpha-frequency

The instantaneous alpha-frequency was extracted for each data point during a one-second period preceding the onset of the first stimulus using the method described by Cohen (2014). First, one-second long epochs were created immediately preceding the onset of the first stimulus. To avoid edge artefacts, each epoch was reflected on both sides; it was flipped horizontally and concatenated to the beginning and end of the original epoch. The data was filtered in the time domain using a plateau-shaped 8-to-13 Hz band-pass filter with 15 % transition zones and a filter order of 896 points. The analytic signal was computed using the Hilbert transform. The phase-angle time series was then unwrapped and its first temporal derivative multiplied by the sampling rate and divided by 2 pi in order to obtain instantaneous frequency in Hz. Noise in the data can cause small jumps in the phase-angle time series (“phase slips”) which in turn produce large artefactual peaks and troughs in instantaneous frequency. To attenuate these, the instantaneous-frequency time-series was median-filtered ten times using ten filter orders ranging from 10 to 400 ms in length, before averaging the ten filtered time series.

Finally, the one-second long time series was divided into 32 time points consisting of 32 samples each (a total of 1024 samples per second). For each trial the average instantaneous frequency over these samples was calculated for each time point. Hence, single-trial instantaneous alpha frequency at 32 time segments and 64 electrodes was entered into the subsequent analysis.

#### Jackknife analysis of the relationships between temporal sensitivity and pre-stimulus EEG power and instantaneous alpha frequency

In order to test whether moment-to-moment fluctuations in pre-stimulus EEG characteristics co-varied with moment-to-moment fluctuations in temporal sensitivity, we implemented a two-level analysis. At the participant level, a single-trial analysis was performed, in which we computed a jackknife Spearman correlation across trials between (i) the JND and EEG power at all time points, frequencies and electrodes and between (ii) the JND and instantaneous alpha frequency at all time points and electrodes. At the group level, these results were subjected to cluster-based permutation tests, to test whether any clusters of data points showed a systematic relationship (i.e. positive or negative correlation) across participants.

##### Single-trial jackknife correlations at the participant level

Since our measure of temporal sensitivity, the JND, is an ensemble metric which cannot be obtained at the single-trial level, the application of jackknife correlations was required. In jackknife correlations, the metrics of interest (EEG parameters and the JND in our case) are computed iteratively over all trials while one trial is left out on each iteration (and reinserted at the next). This results in variables which contain as many values for the statistic as there are trials. Since the resulting statistic at each trial reflects the effect of that trial being left out, the direction of variance is inverted. Therefore, any correlation of a variable with its jackknife counterpart is −1. Additionally, since the effect is scaled by the number of trials, the variance is compressed. This inversion and compression of variance resulting from the jackknife procedure is illustrated in supplementary figure S1. Please note that because both the behavioural data (the JND) and the electrophysiological data (power and instantaneous alpha frequency) were subjected to the jackknife procedure, the correct sign of the resulting correlations was restored. This method enabled us to test the relationship between fluctuations in EEG parameters and temporal sensitivity on a short, trial-to-trial time scale. For the mathematical proof of the equivalence of the conventional and jackknife correlations we refer to Richter et al. (2015). For a detailed explanation of how to apply this procedure to link psychophysical data with EEG data, see (Benwell, et al., 2018). We used Spearman’s rho (*r*_s_) to correlate both EEG power and instantaneous alpha frequency with the JND across trials. For EEG power, this procedure was repeated at all electrodes, frequencies and time points resulting in a 64 ⨯23 ⨯ 25 matrix of *r*_*s*_’s per participant. For instantaneous alpha frequency, we repeated the procedure for all time points and electrodes resulting in a matrix of 64 ⨯ 32 *r*_*s*_’s per participant. Importantly, we controlled for possible non-stationarities in power and frequency over the course of the experiment by partializing out trial order (Pearson, 1915). This precluded the possibility that a spurious correlation would arise due to co-occurring but unrelated EEG and behavioural non-stationarities over the course of the experiment (Benwell et al., 2019).

##### Group-level analysis

Subsequently, we tested whether any of the correlations obtained at the participant level showed a systematic deviation from zero across participants. Dependent sample t-tests against 0 were performed on the Spearman *r*_*s*_’s at each data point across participants. To control for multiple comparisons, a cluster-based permutation-testing routine developed by Maris and Oostenveld (2007) was implemented. This was done separately for correlations of behaviour with EEG power and instantaneous alpha frequency. All data points were selected for which the *t*-value had a probability lower than 5% of having occurred by chance. These were then clustered based on adjacency (at least one channel adjacent to a significant data point had to be significant for cluster inclusion) in the temporal, spectral or spatial domain for EEG power and in the temporal and spatial domain for instantaneous alpha frequency. For the EEG power analysis, this procedure was done separately for positive and for negative *t*-values (two-tailed test). For the instantaneous alpha frequency analysis, a one-tailed test was employed and only negative *t*-values were clustered (we hypothesized that higher instantaneous alpha frequency would be associated with a smaller JND based on Cecere et al. (2015) and Keil and Senkowski (2017). For each cluster, the sum of *t*-values was then calculated and the maximum of these cluster-level statistics was taken. To create a reference distribution against which to test the value of this cluster-level statistic, 1000 permutations of the data were conducted using the “ft_statistics_montecarlo” function from the fieldtrip toolbox (Oostenveld et al., 2011). Each iteration yielded a maximum cluster level statistic and over iterations a null distribution of maximum cluster level values was constructed. The *P*-value of the effect was then estimated as the proportion of elements in the null distribution exceeding the observed maximum cluster-level test statistic.

### Electrophysiological analyses of individual differences: Co-variations of JND and pre-stimulus EEG across participants

To test whether individual-peak alpha-frequency and individual alpha power could predict temporal sensitivity across individuals, we calculated individual-peak alpha-frequency during the pre-stimulus period for each participant. Then, we calculated alpha power at each participant’s individual-peak alpha-frequency. Both these measures were then correlated with the JND across participants using a Spearman correlation. We corrected for multiple comparisons with a Bonferroni correction. Seven participants who did not exhibit a clear alpha peak were excluded from this analysis (see below for details).

For each participant, the presence of an individual alpha peak was estimated from the following posterior electrodes; TP7, CP5, CP3, CP1, CPz, CP2, CP4, CP6, TP8, P9, P7, P5, P3, P1, Pz, P2, P4, P6, P8, P10, PO7, PO3, POz, PO4, PO8, O1, Oz, O2, Iz. The frequency range of interest was 8-13 Hz. The same data epochs as for the within participant analysis described above were included. Each segment was multiplied with a Hanning window and the data were zero padded to obtain a frequency resolution of 0.25 Hz. In order to determine individual-peak alpha-frequency, an automated estimation routine was utilized developed by Corcoran et al. (2018). This provided us with an objective and replicable method to pick the electrode from which the individual-peak alpha-frequency was inferred, and to include only participants who exhibited a clear alpha peak. The method is described in detail in Corcoran et al. (2018). First, the power spectrum is smoothed using a Savitzky-Golay filter. This is a least-squares polynomial curve-fitting procedure originally developed in chemistry to detect spectral peaks amidst noisy conditions (Savitzky & Golay, 1964). The first- and second-order derivatives of the fitted function are then analyzed for evidence of a peak in the alpha band (8-13 Hz). A peak was considered to be valid when the highest power value was at least 1 standard deviation from the mean and was at least 20 % higher than other peaks in the same spectrum. As a measure of the quality of each peak, the area under the peak was defined and divided by its frequency span. Only those participants for whom at least three electrodes yielded a peak were retained in order to exclude those where the detected alpha peak may be spurious. Thirty-three participants satisfied this condition. The electrode with the largest value was chosen to provide the individual-peak alpha-frequency estimate for that participant. Individual alpha power was determined as power at this individual-peak alpha-frequency.

Peaks were detected at the following electrodes (1 per participant; see figure 5A); 1xP3, 1xP5, 5xPO7, 2xPO3, 1xO1, 3xOz, 8xPOz, 1xPz, 1xCPz, 1xP6, 3xPO8, 4xPO4, 2xO2.

### Follow-up analysis of EEG alpha power and instantaneous alpha frequency

To tease apart the power- and instantaneous alpha-frequency effects in the time-frequency analysis (see results section), we conducted a follow-up analysis of activity in the alpha band exclusively. First, we tested whether there was a differential contribution of upper versus lower band alpha-power to the power effect. Then, we tested whether such an asymmetry could explain our instantaneous alpha frequency findings. Additionally, we correlated the power effect with the instantaneous alpha frequency effect across participants. Finally, we correlated the magnitude of the asymmetry of the power effect, with the instantaneous alpha frequency effect across participants.

#### Time-Frequency analysis (alpha band) centered on individual peak alpha frequency

In order to examine contributions of the lower-versus upper alpha bands separately, we accounted for individual differences in alpha peak frequency and aligned participant’s spectra on their individual alpha peak frequency as follows. Power was obtained as described above for the time-frequency analysis, but with a higher frequency resolution of 1 Hz. Single-trial oscillatory power was obtained at 25 time points, from −750 ms to −250 ms in 20 ms steps and across 10 frequencies ranging from 6 to 15 Hz for all 64 electrodes. Average spectra were then calculated for each participant over the electrodes that were present in both the power and instantaneous alpha frequency clusters (CP3, P3, PO7, O1, POz, O2, PO4, P4; see results section, figures 3B, 4B, 7D). The individual alpha peak was determined for this analysis as the frequency with maximum power averaged over time-points in the range from 8 to 13 Hz, with the condition that the spectrum (from 6 to 15 Hz) did not decrease monotonically (which was the case in 38 out of 40 participants). Two participants had spectra that decreased monotonically and were hence excluded from this follow-up analysis. For each individual we constructed the alpha power spectrum by taking their individual alpha peak and including the frequencies up to two Hz above and below it. This resulted in a time-frequency spectrum of 5 frequency points x 25 time points for each of the 8 electrodes. Then, as we did in the original time-frequency analysis, we correlated power with the JND across trials for each channel and time-frequency point, while controlling for trial order. Since the data points we chose for this follow-up analysis were based on inclusion in a significant cluster, we did not perform the cluster-based permutation test again, to avoid double-dipping (Kriegeskorte et al., 2009). We report whether the uncorrected *P*-value at each channel-time-frequency point was below 0.05 for illustrative purposes.

**Figure 3.**
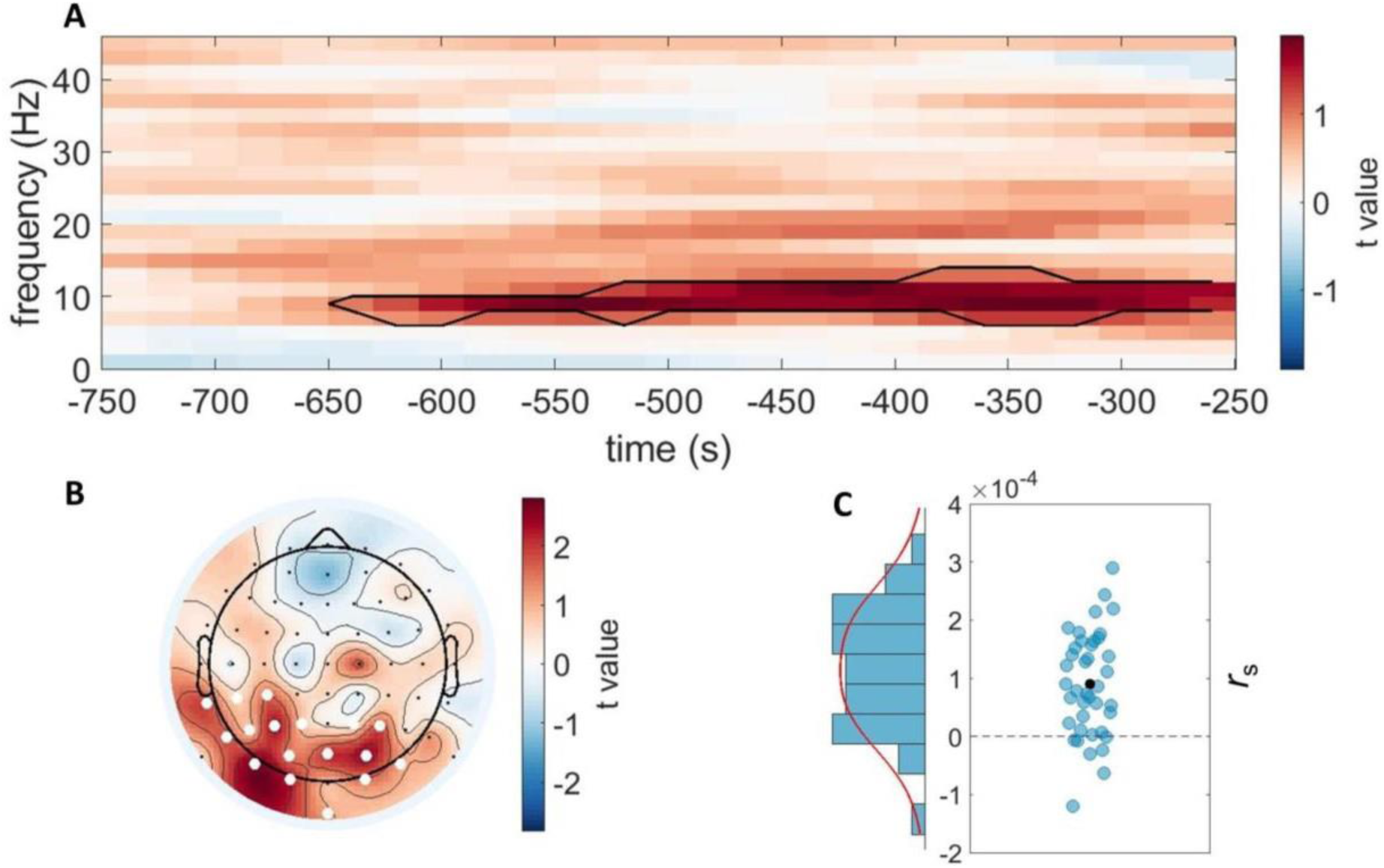
EEG oscillatory power predicts the temporal sensitivity of audio-visual perception. **(A)** Time-frequency representation of t-values averaged over all electrodes included in the significant cluster. Onset of the first stimulus (auditory or visual) is at 0 seconds. One positive cluster survived multiple comparison correction and is outlined in black. The positive t-values (in red) indicate that higher power was accompanied by a higher JND (worse temporal sensitivity). No negative clusters were observed. **(B)** Topographical representation of the t-values averaged over all time-frequency points included in the significant cluster. Electrodes that were included in the significant cluster are highlighted in white. **(C)** Average correlation coefficients for each participant between EEG power in the cluster and the JND. Each blue dot represents one participant. The black dot indicates the group mean, and the black dotted line indicates a correlation of 0.

#### Instantaneous alpha frequency contribution to JND: Controlling for alpha-power

Next, we tested whether instantaneous alpha frequency had any independent predictive value with regard to the JND. In order to relate the instantaneous alpha frequency results with the power results, since our power analysis only included data from 750 to 250 ms pre-stimulus, we cut the instantaneous frequency data to match this time window and time points. Then, as we did in the original time-frequency analysis described above, we calculated Spearman’s ϱ between instantaneous alpha frequency and the JND across trials for each time point over the electrodes that were present in both the power and instantaneous alpha frequency clusters (CP3, P3, PO7, O1, POz, O2, PO4, P4; see results section, figures 3B, 4B, 7D). Subsequently, we repeated the analysis five more times, each time controlling for alpha power at each one of the five individualized alpha frequencies (peak alpha - 2 Hz, peak alpha - 1 Hz, peak alpha, peak alpha + 1 Hz, peak alpha +2 Hz). We controlled for alpha power by computing partial correlations (Pearson, 1915) between the JND and instantaneous alpha frequency, while partializing out alpha power (and trial order, as we did in all analyses). As with the alpha-power analysis described above, the two participants for whom the alpha spectra decreased monotonically were excluded from this analysis. Again, we did not perform the cluster-based permutation test, but report whether the uncorrected *P*-value at each channel-time-frequency point was below 0.05.

#### Correlation analysis across participants

If the power- and instantaneous alpha frequency effects are related within participants, then we might expect them to be related across participants as well. First, we tested whether the effects we found in the original analyses were related across participants. For each participant we took the correlations between EEG power and the JND and averaged them over all data points in the significant cluster (as presented in fig. 3C). We did the same for the correlations between instantaneous alpha-frequency and the JND (as presented in fig. 4C). Then, using a Spearman correlation, we correlated these ϱ values across participants. Second, we turned to the results from the follow-up analysis, involving all time-points from 750 to 250 ms before stimulus presentation, and used only those eight electrodes that were present in both the power and instantaneous alpha frequency clusters from the original analyses. We tested whether it might be the upper/lower band asymmetry of the alpha power – JND relation that predicted the instantaneous alpha frequency – JND relation. We quantified this asymmetry by averaging the correlation coefficients of the alpha power – JND relation over the upper alpha band (individual-peak alpha-frequency + 1Hz and individual-peak alpha-frequency + 2Hz) and over the lower alpha band (individual-peak alpha-frequency - 1Hz and individual-peak alpha-frequency - 2Hz) separately and subtracting the average of the upper alpha band from the average for the lower alpha band. Then, using a Spearman correlation, we correlated this alpha asymmetry with the instantaneous alpha frequency – JND relation (uncontrolled for alpha power) across participants.

**Figure 4.**
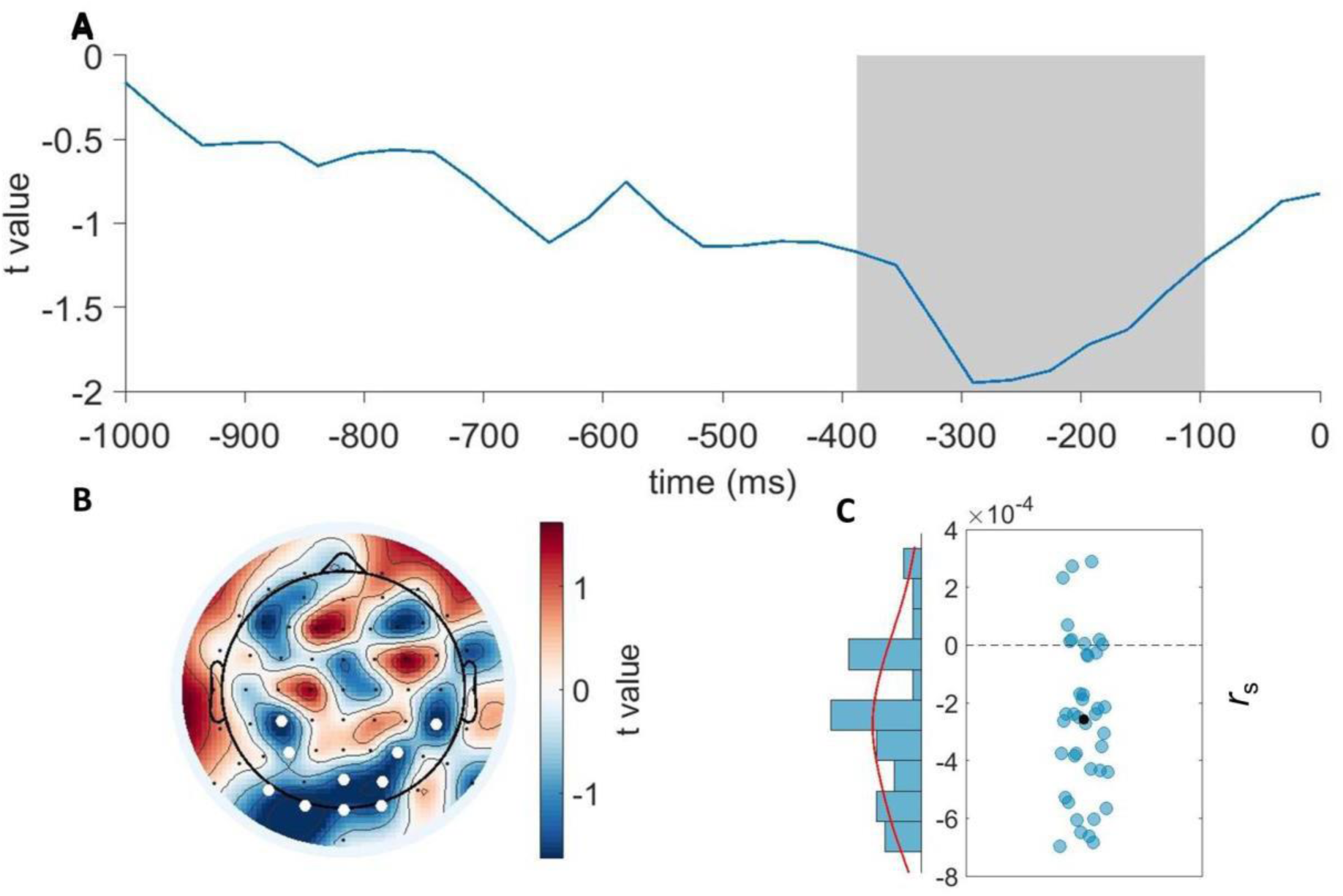
EEG instantaneous alpha frequency predicts the temporal sensitivity of audio-visual perception. **(A)** t-values over time, averaged over all electrodes included in the cluster. Onset of the first stimulus (auditory or visual) is at 0 seconds. Negative t-values indicate that higher instantaneous frequency is accompanied by a lower JND (higher temporal sensitivity). One negative cluster survived multiple comparison correction and is marked with a grey box. **(B)** Topographical representation of the t-values averaged over all time points included in the cluster. Electrodes that were included in the cluster are highlighted in white. **(C)** Average correlation coefficients for each participant between instantaneous alpha frequency in the cluster and the JND. Each blue dot represents one participant. The black dot indicates the group mean and the dotted line indicates a correlation of 0.

**Figure 5.**
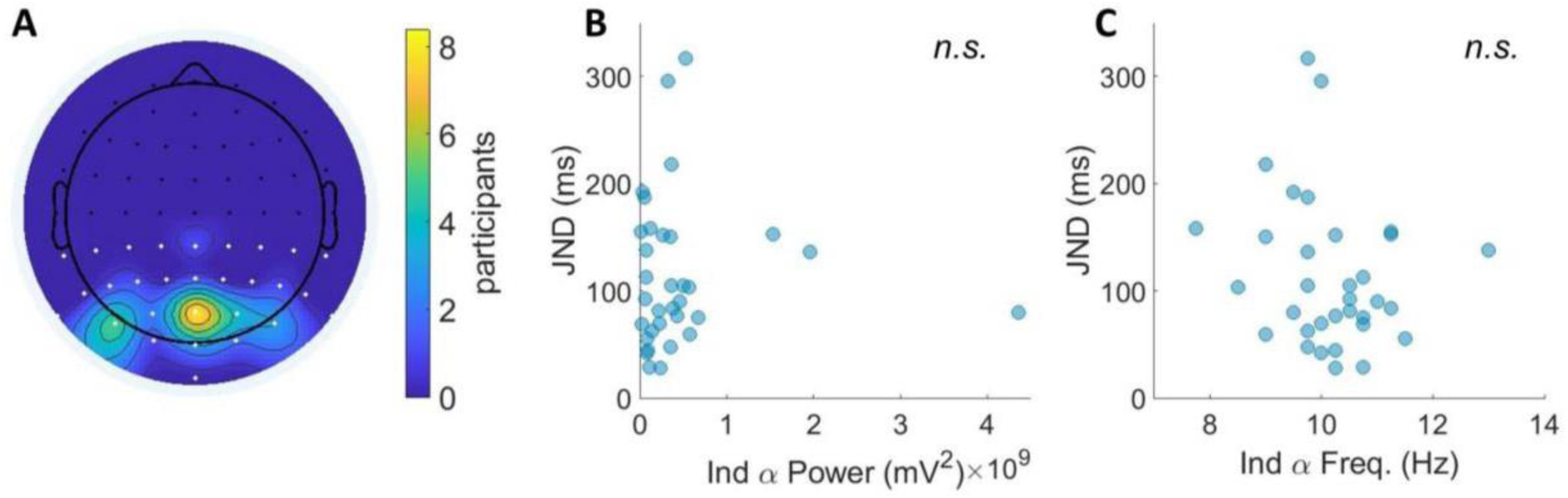
Individual alpha power and individual alpha frequency did not predict temporal sensitivity of audio-visual perception across participants. **(A)** Heat map showing the number of participants for which each electrode was selected to measure individual-peak alpha-frequency and individual-peak alpha-power. The electrodes that were entered into the peak-identification algorithm are marked in white. **(B)** No systematic relationship was found between individual-peak alpha-power and JND. **(C)** No systematic relationship was found between individual-peak alpha-frequency and JND.

## Results

### Behavioural results

Participants completed an audio-visual TOJ task. They were presented with a beep and a flash at varying SOAs, and were asked to indicate which stimulus had been presented first (Figure 1). A psychometric function was then fitted to the proportion of “flash-first” responses. Figure 2A shows the fitted functions for each participant as well as a function fit to the average data of all participants (in black). As an index of temporal audio-visual discrimination sensitivity, - our primary measure of interest, - we derived the JND from this function. Across all participants the mean JND was 110 ms (st. dev. of 64 ms), which is typical of the large individual differences previously observed in such paradigms (e.g. Stevenson et al., 2012; see figure 2B). Additionally, we derived the point of subjective simultaneity (PSS). On average the PSS was −21 ms (audio-leading). However, this deviation from 0 was not statistically significant (the 95% confidence interval ranged from −44.41 ms to 3.11 ms). See Figure 2B and C for individual data points.

### EEG results

#### Pre-stimulus alpha power predicts temporal sensitivity

We tested if the power of spontaneous fluctuations during the pre-stimulus interval predicted the temporal sensitivity of audio-visual perception on a trial-by-trial basis. Figure 3A shows the strength and direction of the relationship between EEG power and the JND in timefrequency space. One significant positive cluster (*P* = .018) was present in the alpha frequency range from 650 to 250 ms preceding stimulus onset. The significant cluster was restricted to posterior electrodes (see figure 3B). The results indicate that higher pre-stimulus power was associated with higher JND values and hence lower sensitivity. Figure 3C shows for each participant the correlation between power and the JND averaged over the points in the cluster.

For 33 out of 40 participants (82.5%), higher power was accompanied by worse temporal sensitivity. This was not a spurious finding caused by coexisting, but independent, changes in alpha power and JND over the course of the experiment (due to fatigue, boredom and/or decreased motivation; see Benwell et al., 2019), as the analysis controlled for trial order. Therefore, these data suggest a functional role of ongoing alpha oscillatory power in the temporal sensitivity of audio-visual perception on a short, trial-to-trial time scale.

#### Pre-stimulus instantaneous alpha frequency predicts temporal sensitivity

We also tested whether instantaneous frequency of alpha oscillations in the pre-stimulus period (1000 ms window) co-varied with temporal sensitivity of audio-visual perception on a trial-by-trial basis. Based on previous findings (Cecere et al., 2015; Keil & Senkowski, 2017; Samaha & Postle, 2015; Wutz et al., 2018), we expected higher instantaneous alpha frequency to predict higher temporal resolution, and hence a negative relationship with JND (higher frequency – smaller JND). Figure 4A shows the strength of the relationship between alpha instantaneous frequency and the JND over the pre-stimulus interval in the only significant (negative) cluster. The time segment where higher alpha instantaneous frequency significantly predicted a lower JND (and thus better temporal sensitivity) is marked with a grey box (*P* = .024). This cluster was present from 387 ms to 97 ms before stimulus onset. Only posterior electrodes were included in this cluster (see Fig. 4B). The cluster strongly overlapped in time and space with the pre-stimulus power cluster (cf. Fig 3B). Figure 4C shows for each participant the correlation between alpha instantaneous frequency and the JND averaged over the points in the cluster. For 31 out of 40 participants (77.5%), higher instantaneous alpha frequency was accompanied by better temporal sensitivity. Therefore, these data are also in line with the existence of a functional role of alpha oscillatory frequency in the temporal sensitivity of audio-visual perception on a short, trial-by-trial time-scale.

#### Neither individual peak-alpha frequency nor individual peak-alpha power predicts individual differences in temporal sensitivity

So far, we found that trial-by-trial variations in both alpha power and instantaneous alpha frequency predict the temporal sensitivity of multisensory perception at the group level. Next, we tested whether alpha power and frequency could also predict performance across individuals, and thereby help explain the large inter-individual variability in JND’s typically found in such paradigms (e.g. Stevenson et al., 2012). We calculated individual peak alpha frequency during the pre-stimulus period averaged over all trials for each participant. We then calculated mean alpha power at this frequency (individual-peak alpha-power) and correlated both measures with the individual JNDs across participants using a Spearman correlation. The results did not reveal any significant correlation across participants, even when we only included participants with a clear alpha peak (*n* = 33) based on a conservative peak-finding algorithm (see methods). Figure 5 shows that neither individual peak-alpha frequency (Fig. 5B; *r*_*s 31*_ = −0.21, *P* = 0.12, one-tailed) nor individual peak-alpha power (Fig. 5C; *r*_*s 31*_ = 0.03, *P* = 0.85, two-tailed) were predictive of an individual’s JND.

#### Alpha frequency and power effects on temporal sensitivity appear to be driven by the same process

A question that arises is whether the power and frequency effects on JND reflect independent predictors of temporal sensitivity or alternatively are connected, and if so, how? When measuring the instantaneous (alpha) frequency of a scalp-level signal such as the EEG, it is important to keep in mind that it likely includes contributions from multiple oscillatory sources, even when it is bandpass filtered around the frequency band of interest (Benwell et al., 2019; Clayton et al., 2017). Hence, power and frequency predictors of perception in the alpha band may originate from different processes, even if recorded from overlapping sites. On the other hand, the instantaneous frequency and power of a signal are not independent in many instances (Nelli et al., 2017) as a result of which the links of power and frequency to JND may be connected. One scenario in which these effects may result from the same underlying process is if the driving force of both effects were an asymmetric power variation in the alpha band (see figure 6). More specifically, a selective variation (increase or decrease) in lower alpha power would result in an opposite variation (decrease or increase) in instantaneous alpha frequency (see figure 6), because the instantaneous frequency of the summated scalp signal will gravitate towards the frequency of the oscillatory source with the highest power. This pattern would be in line with our results; the instantaneous alpha frequency relationship with the JND is negative, whereas the relationship of power with the JND is positive. However, if the power effect in our data were mostly driven by upper rather than lower band alpha signals, then this could not explain the negative relation we found between instantaneous alpha frequency and the JND, nor could an effect of power at peak alpha frequency.

**Figure 6.**
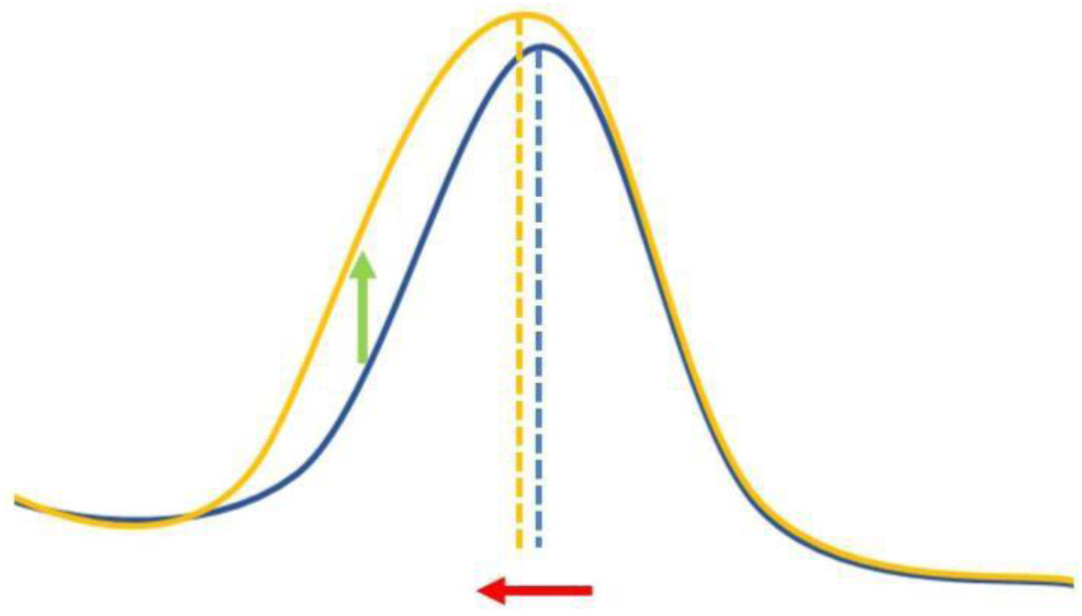
Toy model depicting the scenario where a variation of lower-band alpha power could lead to a concurrent modulation of instantaneous alpha frequency. The power spectrum in yellow shows an increase in mostly lower-alpha power compared to the blue spectrum. This increase in lower-alpha power pulls the instantaneous alpha frequency measurement of the signal towards the lower alpha range, where more power is now present. Conversely, a selective decrease in lower-band alpha power would pull the alpha frequency towards the higher alpha range (not shown).

A reanalysis of our data centered on the alpha band was in support of the power and frequency effects resulting from the same process, as the strongest effect of power was observed in the lower alpha band (figure 7). Power at individual alpha - 1 Hz showed the most pronounced relationship with the JND, as reflected in higher average *t*-values and a greater number of electrodes and data points showing an uncorrected *P*-value below .05 (see figure 7A). It is of course still possible that the instantaneous frequency effect is only partially due to fluctuations in lower-alpha power, and that instantaneous alpha frequency also has an independent relation with the JND. To test this, we correlated instantaneous alpha frequency and the JND across trials again. Crucially, we now controlled for alpha power. For comparison, figure 7B shows the *t*-values averaged over all 8 electrodes, for the instantaneous alpha frequency – JND relation without controlling for alpha power. Figure 7C shows the *t*-values averaged over all 8 electrodes, for the instantaneous alpha frequency – JND relation with each row representing the results when controlling for alpha power at each one of the five individualized alpha frequencies. The *t*-values reflecting the relationship between instantaneous alpha frequency and the JND, as well as the number of electrodes and data points showing an uncorrected *P*-value below .05, were markedly reduced when controlling for power at individual alpha – 1 Hz and – 2 Hz, with the strongest reduction at −1 Hz, where the strongest effect for alpha power was found. The inverse was true when controlling for power at individual alpha + 1 Hz and + 2 Hz; here the *t*-values reflecting the relationship between instantaneous alpha frequency and the JND, as well as the number of electrodes and data points showing an uncorrected *P*-value below 0.05 were increased. This pattern of results is consistent with the idea that the analyses of power and instantaneous alpha frequency in relation to the JND reveal the same process. When power is higher at *lower* frequencies, the resulting measure of instantaneous alpha frequency will be lower. Both higher power and lower instantaneous frequency are related to a higher JND. Controlling for power in the lower frequencies (individual alpha – 1 Hz and – 2 Hz) when correlating instantaneous alpha frequency with the JND will therefore weaken the correlation coefficient. Conversely, when power is higher at *higher* frequencies, the resulting measure of instantaneous alpha frequency will also be higher. Whereas higher power is related to a higher JND, higher instantaneous alpha frequency is related to a lower JND. Therefore, controlling for power in the higher frequencies (individual alpha + 1 Hz and + 2 Hz) when correlating instantaneous alpha frequency with the JND will strengthen the correlation coefficient. Finally, we tested whether we could find a similar overlap between the alpha power – JND relationship and the alpha instantaneous frequency – JND relationship across participants. The relationship between the average magnitude of the alpha power – JND correlation shown in fig. 3C, and the instantaneous alpha frequency - JND correlation shown in fig. 4C across participants was not significant (*r*_*s 38*_ = -.18, *p* = .27; see fig. 7E). We also tested whether it might be the upper/lower band asymmetry of the alpha power – JND relation that predicted the instantaneous alpha frequency – JND relation across participants. This asymmetry was calculated by averaging the correlation coefficients of the alpha power – JND relation over the upper alpha band (individual alpha + 1 Hz and + 2 Hz) and over the lower alpha band (individual alpha – 1 Hz and – 2 Hz) separately and then subtracting the average of the upper alpha band from the average for the lower alpha band. The magnitude of the asymmetry of the alpha power – JND relation was highly predictive of the correlation between instantaneous alpha frequency and the JND (*r*_*s 36*_ = -.79, *P =* .000000013; see fig. 7F). In other words, a stronger correlation between alpha power per sé and the JND did not predict the correlation between instantaneous alpha frequency and the JND, but the asymmetry of the correlation between high and low alpha power and the JND did. The results of this follow-up analysis suggest that a modulation of power in the lower-alpha band predicts the temporal resolution of audiovisual integration in our paradigm and that this effect is reflected in correlations (of opposite sign) between both alpha power and instantaneous alpha frequency and the JND.

**Figure 7.**
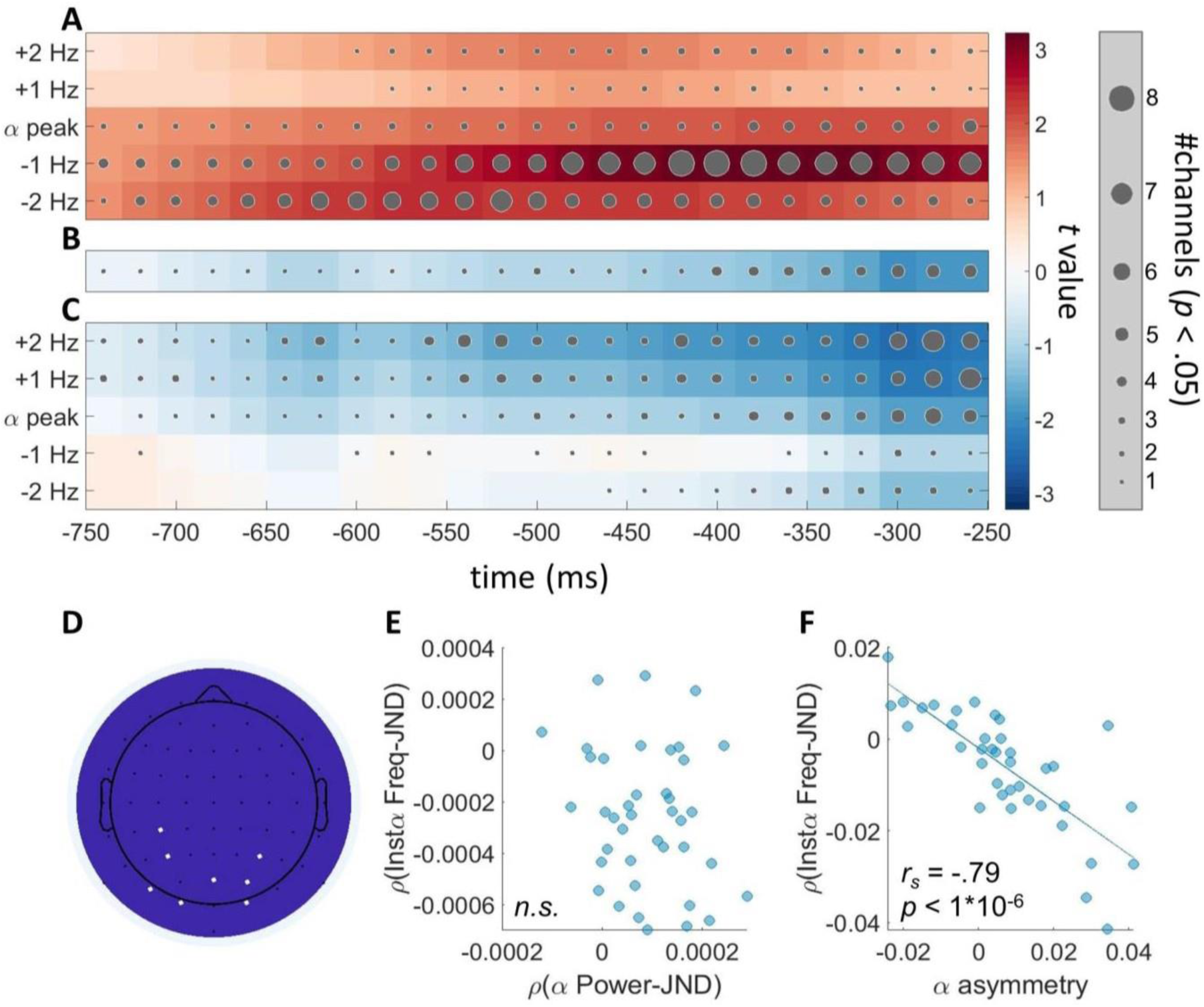
The predictive value of instantaneous alpha frequency for the temporal sensitivity of audio-visual perception is linked to fluctuations in lower-alpha power. (**A**) Time-frequency representation of t-values centred on the individual alpha peak frequency for the relationship between alpha power and the JND, averaged over all electrodes included in both the power and instantaneous alpha frequency clusters depicted in figures 3B and 4B (see panel D). Positive t-values (in red) indicate that higher power is accompanied by a higher JND (worse temporal sensitivity). The size of the dots indicates the number of electrodes out of 8 with an uncorrected p-value < .05 at that time point. (**B**) Time-frequency representation of t-values for the relationship between instantaneous alpha frequency and the JND, uncontrolled for alpha power, averaged over all electrodes included in both the power and instantaneous alpha frequency clusters depicted in figures 3B and 4B (see panel D). Negative t-values (in blue) indicate that higher instantaneous frequency is accompanied by a lower JND (higher temporal sensitivity). The size of the dots indicates the number of electrodes out of 8 with an uncorrected p-value < .05 at that time point. (**C**) Time-frequency representation of t-values for the relationship between instantaneous alpha frequency and the JND, controlled for alpha power, averaged over all electrodes included in both the power and instantaneous alpha frequency clusters depicted in figures 3B and 4B (see panel D). Each row represents one of 5 individualized alpha frequencies being controlled for. Negative t-values (in blue) indicate that higher instantaneous frequency is accompanied by a lower JND (higher temporal sensitivity). The size of the dots indicates the number of electrodes out of 8 with an uncorrected p-value < .05 at that time point. (**D**) The electrodes on which the analyses in panels A,B,C and F were based (marked in white). (**E**) Spearman correlation between the average magnitude of the alpha power – JND correlation shown in fig. 3C, and the instantaneous alpha frequency - JND correlation shown in fig. 4C across participants (**F**) Spearman correlation between the magnitude of the upper/lower band asymmetry of the alpha power – JND relationship shown in fig. 7A, and the instantaneous alpha frequency - JND correlation shown in fig. 7B.

## Discussion

We used a temporal order judgement (TOJ) task to examine the role of spontaneous, ongoing EEG oscillations in the temporal sensitivity of audiovisual integration. Pre-stimulus power at a wide range of frequencies was tested, and we found that alpha power predicted performance at the single-trial level. Lower power in this frequency band (8-13 Hz) predicted better temporal sensitivity. The single-trial instantaneous alpha frequency was also measured and higher instantaneous alpha frequency was found to predict better temporal sensitivity. At first glance, the latter result seemed to support an account of audiovisual temporal integration in terms of the theory of perceptual cycles (VanRullen, 2016). A follow-up analysis revealed, however, that the correlation between instantaneous alpha frequency and the JND could be explained by power modulations in the lower alpha band (below individual alpha peak).

These results provide novel insights into the neural basis of the temporal resolution of multisensory integration. We show that not only task conditions (Stevenson & Wallace, 2013; van Eijk et al., 2008) and individual differences (Stevenson et al., 2012; Wallace & Stevenson, 2014) affect the temporal sensitivity of audiovisual integration, but that spontaneous alpha oscillations do so as well. In showing this, we extend existing evidence that higher alpha power is indicative of a tendency towards temporal integration (Bastiaansen et al., 2020; Baumgarten et al., 2016; Leonardelli et al., 2015; Peterson & Voytek, 2018). Furthermore, our pattern of results highlights the need for caution when interpreting analyses involving instantaneous alpha frequency, since asymmetric variations of power in the upper/lower alpha band, such as we observed here, can also be reflected in a (possibly epiphenomenal) relationship between instantaneous alpha frequency and behaviour. Such asymmetric power shifts may turn out to be common, as evidence for the existence of multiple, variable alpha rhythms is mounting. These rhythms are believed to originate from both cortical and sub-cortical sources that together give rise to the rhythm measured at the scalp (Benwell et al., 2019; Clayton et al., 2017; Klimesch et al., 1996).

### Decreased alpha power predicts increased temporal sensitivity

We found that lower alpha power predicted better temporal sensitivity in an audiovisual TOJ. This effect was driven mostly by power in the lower-alpha band as compared to power in the mid- or upper-alpha band and the effect was most pronounced over occipito-parietal electrodes. There is some evidence that activity in the lower- and mid-alpha band reflects expectancy and alertness, with power decreasing as alertness increases, and that upper alpha band activity reflects cognitive performance in certain tasks, with alpha power rising as performance improves (Klimesch et al., 1998). Even though this evidence is limited and comes mostly from studies into memory maintenance, the idea that alpha band activity reflects expectancy and alertness fits well with evidence that pre-stimulus alpha oscillations index excitability of the cortex, with higher alpha power indicating lower excitability (Romei et al., 2008; Sauseng et al., 2009). In studies where participants are asked to detect a weak stimulus, lower pre-stimulus alpha power commonly leads to higher detection rates (as reviewed in Iemi et al., 2017). Notably, this is the case whether the stimulus is real or illusory. Thus, lower pre-stimulus alpha power does not necessarily lead to more accurate perception (e.g. Benwell, et al., 2018; Lange et al., 2013). Iemi et al. (2017) addressed this issue with signal detection theory. They hypothesized that if decreased alpha power indicates increased baseline excitability, not only the signal but also the noise would elicit a larger response. This would lead to more hits, but also to more false alarms, thereby shifting the criterion towards the more liberal side, but leaving sensitivity unchanged. Indeed, they found that in a near-threshold visual stimulus detection task, decreased alpha power made observers more likely to report the presence of a stimulus, whether the stimulus was present or not. In a discrimination task, they found that alpha power did not affect performance, in accordance with the idea that perceptual bias, but not sensitivity is affected by alpha oscillations. Other studies on visual perceptual discrimination sensitivity have also shown this measure to be unaffected by alpha power shifts (Bays et al., 2015; Benwell et al., 2017, 2018; Lou et al., 2014; Wutz et al., 2014). Our data do not mirror these results. We found that pre-stimulus alpha power did predict discrimination sensitivity. Similar results have been reported by Leonardelli et al. (2015) who presented participants with an audio-tactile pair of above-threshold stimuli with variable SOA’s while recording the magneto-encephalogram. When comparing brain activity between identically timed pairs with different perceptual outcomes they found that on trials where participants perceived one integrated audio-tactile stimulus, pre-stimulus alpha power had been higher compared to trials where participants perceived the stimuli as separate. On a comparable note, (Baumgarten et al., 2016) presented participants with one or two short, above-threshold tactile stimuli. When the time between the stimuli was such that the percept varied from 1 to 2 on a trial-by-trial basis, decreased pre-stimulus alpha power predicted veridical perception of 2 stimuli. In other words, in a tactile temporal discrimination task (Baumgarten et al., 2016), in an audio-tactile temporal discrimination task (Leonardelli et al., 2015), and in our audiovisual temporal discrimination task, lower alpha power predicted higher temporal sensitivity and segregated perception, while higher alpha power predicted lower temporal sensitivity and integrated perception. These studies differ in at least three characteristics from the visual discrimination tasks where alpha power did not affect perceptual sensitivity (Bays et al., 2015; Hanslmayr et al., 2007; Wutz et al., 2014; Benwell et al., 2017, 2018). First, they are not unisensory visual tasks, but either multisensory (our study and Leonardelli et al., 2015) or do not involve the visual modality at all (Baumgarten et al., 2016). Second, above-threshold stimuli were presented instead of near-threshold stimuli. And third, they involve temporal discrimination, whereas the mentioned visual tasks involve discrimination based on visual features such as orientation (Bays et al., 2015), identity (Hanslmayr et al., 2007) or numerosity (Wutz et al., 2014). Temporal discrimination differs in a fundamental manner from these visual feature criteria in that it requires perception to be updated on a short time-scale. There is evidence that alpha power actually promotes stability as opposed to the flexibility required for this fast updating. For example, when viewing a Necker cube, perception spontaneously alternates between two rivalling perceptual interpretations (Necker, 1832). In this paradigm, higher alpha power correlates with a longer duration of each of the rivalling percepts and thus higher perceptual stability (Piantoni et al., 2017) and reductions in alpha power predict an impending switch from one percept to the other (Strüber & Herrmann, 2002). Based on these studies, (Piantoni et al., 2017) proposed that alpha oscillations do not purely inhibit cortical activity but stabilize the current configuration of neuronal activity and its corresponding perceptual interpretation. Despite the lack of spatial specificity of the EEG, it is interesting to note that the relation between alpha power and temporal sensitivity in our experiment is most pronounced over occipito-parietal areas, which mirrors the topographies of the relationship between alpha power and perceptual stability in Piantoni et al.’s (2017) study. In temporal discrimination tasks such as ours, higher excitability would lead to an improvement of sensitivity due to a greater perceptual flexibility to adapt to new information on short time-scales.

Studies using other temporal discrimination paradigms have produced results that are in line with ours. For example, (van Viegen et al., 2017) presented participants with a tone and then after 1 or 1.5 seconds a flash. They found that the tone always elicited alpha and beta suppression over parietal and occipital electrodes, but that the long intervals were more likely to be incorrectly perceived as short intervals when alpha and beta power were less suppressed. They concluded that higher alpha and beta power led to a subjective compression of time, which can also be interpreted as stronger integration over time. And in a multisensory time-estimation task, (van Driel et al., 2014) tested how phase coupling between auditory and visual sensory regions was related to interference effects from one modality to the other. They found that when participants had to judge the duration of a visual target, the duration of an auditory distractor interfered more in those participants with stronger alpha phase coupling between auditory and visually responsive electrodes. As in our study, stronger alpha synchronization was indicative of cross-modal temporal integration. Taken together, the evidence suggests that when excitability is low, and alpha synchronization is high, the cortex leans towards temporally integrated perception, and that when excitability is high, and alpha synchronization is low, the cortex leans towards temporally segregated perception.

### Increased alpha frequency does not independently predict increased temporal sensitivity

Previous studies suggest that the length of the cycle of alpha oscillations determines the length of time over which stimuli are integrated (Cecere et al., 2015; Keil & Senkowski, 2017; Ronconi et al., 2018; Samaha & Postle, 2015). These findings support the idea of discrete windows of perception or perceptual cycles (VanRullen, 2016). In this study, we attempted to replicate such results using multisensory, supra-threshold stimuli. Although we found the expected pattern, with higher alpha frequency predicting better temporal sensitivity on a trial-by-trial basis, a follow-up analysis revealed that this effect was linked to an asymmetry in alpha-power modulations, as described above. There seems to be a single process, expressing itself in both power and instantaneous frequency, that has functional significance for the temporal resolution of integration. One possibility could be that it is the size of the cell assemblies involved in perceiving the stimuli that affects the temporal resolution of perception. A larger cell assembly synchronizing across a larger range oscillates slower and has more power than a smaller, more localized cell assembly (Nunez, 2000; von Stein & Sarnthein, 2000). Multimodal perception requires synchronization across a larger range of the cortex, between sensory cortices, than unimodal perception, which only requires synchronization within sensory cortices (e.g. (van Driel et al., 2014; Von Stein et al., 1999). It follows that on trials where integration occurs (and the temporal order of the stimuli cannot be resolved) we would observe both higher power and lower frequency oscillations. Instead of discrete perceptual cycles, with the length of the cycle being determined by alpha frequency, the instantaneous frequency – JND relation could reflect that larger cell assemblies are indicative of integration across a larger cortical area and produce both higher frequency oscillations and higher power. Therefore, despite our initial positive finding, with results in line with those of Ronconi et al., (2018) and Samaha & Postle (2015), we cannot conclude that the length of the cycle of alpha oscillations determines the length of time over which multisensory, above-threshold stimuli are integrated. One reason for our differing pattern of results could be that whereas the integration of uni-sensory veridical and illusory stimuli occurs mostly in the sensory cortices themselves, activity relevant to the TOJ task occurs at least partially in higher association areas (Binder, 2015; Love et al., 2018; Watkins et al., 2006). Another possibility is that, if Ronconi et al. (2018) and Samaha & Postle (2015) also controlled for alpha power in their analyses, they would find similar results to ours.

Our pattern of results stresses the need to account for individual differences in peak alpha frequency, as well as systematic shifts in both alpha frequency and power over the course of an experiment. Importantly, in light of the increasing popularity of instantaneous frequency as a measure in EEG research (e.g. Babu Henry Samuel et al., 2018; Samaha & Postle, 2015; Shen et al., 2018; Wutz et al., 2018), our results caution against an interpretation of such effects in terms of genuine frequency shifts involving one oscillator, without controlling for the possibility of asymmetric power shifts over multiple oscillators.

### Neither individual peak alpha frequency nor power predicts individual differences in temporal sensitivity

Even though alpha power predicted performance on a trial-by-trial basis, we did not find any relationship between alpha power and temporal sensitivity across participants. Nor did we find a relationship between alpha frequency and temporal sensitivity across participants. This is in contrast to findings from Cecere et al. (2015), Samaha & Postle (2015), and Keil & Senkowski (2017) who found a positive correlation between alpha frequency and temporal sensitivity across participants. Again, an important difference between their tasks and ours is that they both involve some form of visual detection with an implicit temporal factor. In our task, the temporal factor is explicitly probed and a discriminatory response is required. Similar to our results, in the same tactile temporal discrimination task as described above (Baumgarten et al., 2016), this relationship was absent across participants (Baumgarten et al., 2017) despite a within-participant relationship between alpha frequency and temporal sensitivity (Baumgarten et al., 2016). One reason this relationship was absent in our data could be that the TOJ task is a much harder and cognitively demanding task than the sound-induced flash illusion used by Cecere et al. (2015) and Keil & Senkowski (2017). At an individual level, many more factors affect the JND than just the speed and power of oscillations, and might do so more strongly. This is readily apparent when looking at the sizes of the JND exhibited by our participants which ranged from 28 to 316 ms (see fig. 3). It is unlikely that the main factor underlying such a broad range of JND’s could be found in the subtle differences in peak frequency between participants. It might be the case that peak frequency and/or power do matter, but that factors such as task engagement or decision-related processes play a much bigger role, drowning out smaller effects. When conducting analyses within participants, these factors are neutralized, enabling the subtler influence of oscillatory characteristics to come to light.

### Conclusion

In this study we tested whether spontaneous, pre-stimulus EEG activity predicts behavioural performance on an audio-visual temporal order judgement task. We found that lower prestimulus alpha power predicted higher temporal sensitivity on a trial-by-trial basis. Higher pre-stimulus alpha frequency also seemed to predict higher temporal sensitivity, but this effect could be attributed to an upper/lower band asymmetry in the effect of alpha power. This pattern of results encourages careful consideration of asymmetric power effects and individual differences in alpha frequency when interpreting results of instantaneous alpha frequency analyses. We did not find any systematic relationship between individual alpha frequency or individual alpha power and temporal sensitivity across participants. Taken together with previous work, our findings suggest that modulations in alpha power index the brain’s tendency for temporal integration vs. segregation on a trial-by-trial basis.

## Acknowledgements

This work was supported by Ghent University (BOF Grant: B/13469/01) to D.T. and the Fund for Scientific Research-Flanders (FWO-V; grant number V429516N) to R.E.L. We also would like to thank dr. Mike X. Cohen for his inspired teaching of the methods applied in this paper and making his scripts available, and dr. Nicolaas Prins for helping us understand the intricacies of psychometric function fitting.

## Author contributions

R.E.L. designed research, performed research, analyzed data and wrote the article. C.S.Y.B., R.C. and G.T. analyzed data and wrote the article. M.Q. performed research and wrote the article. D.T designed research and wrote the article.

## Data accessibility

Data and analysis scripts can be viewed on the Open Science Framework (OSF) platform by following this link: https://osf.io/vwugm/?view_only=47c362c23d004b96b06030114b5271c9 The project will be made public upon publication of the manuscript.

## Abbreviations

EEG: electroencephalogram
FIR: finite impulse response
ICA: independent component analysis
JND: just noticeable difference
PSS: point of subjective simultaneity
SOA: stimulus onset asynchrony
TOJ: temporal order judgement

---

**Figure S1.**
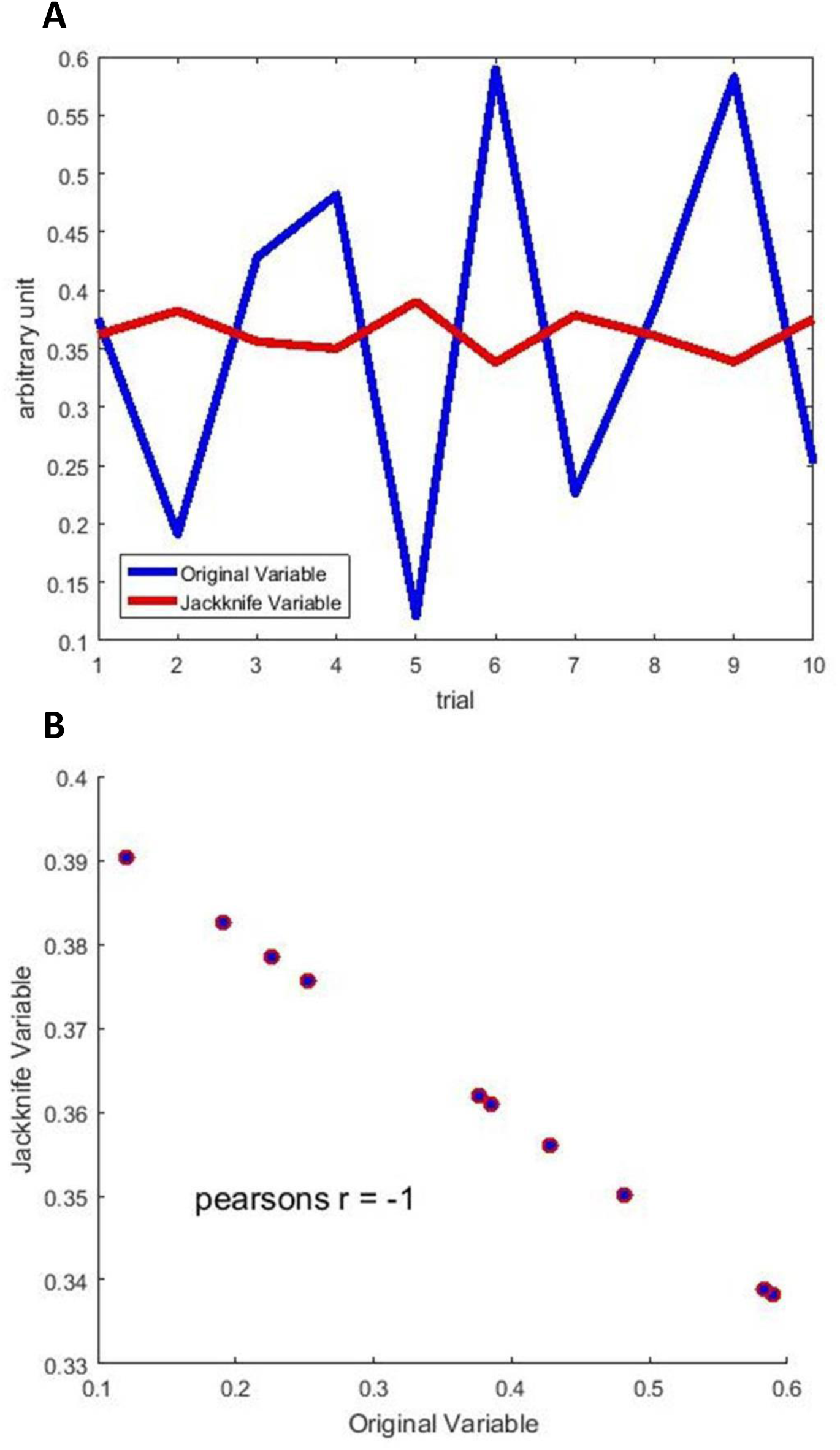
A shows a random variable with 10 values and that same variable after calculating a jackknife statistic (for this example the mean was used). It is apparent that the jackknife variable mirrors the original variable and is scaled down by the number of trials. Figure S1B shows that the original and jackknife variable are perfectly correlated, but that the sign of the correlation is negative.

## References

Babu Henry Samuel, I., Wang, C., Hu, Z., & Ding, M. (2018). The frequency of alpha oscillations: Task-dependent modulation and its functional significance. NeuroImage, 183, 897–906. https://doi.org/10.1016/j.neuroimage.2018.08.063

Bastiaansen, M., Berberyan, H., Stekelenburg, J. J., Schoffelen, J. M., & Vroomen, J. (2020). Are alpha oscillations instrumental in multisensory synchrony perception? Brain Research, 1734, 146744. https://doi.org/10.1016/j.brainres.2020.146744

Baumgarten, T. J., Schnitzler, A., & Lange, J. (2016). Prestimulus Alpha Power Influences Tactile Temporal Perceptual Discrimination and Confidence in Decisions. Cerebral Cortex, 26(3), 891–903. https://doi.org/10.1093/cercor/bhu247

Baumgarten, T. J., Schnitzler, A., & Lange, J. (2017). Beyond the Peak – Tactile Temporal Discrimination Does Not Correlate with Individual Peak Frequencies in Somatosensory Cortex. Frontiers in Psychology, 8. https://doi.org/10.3389/fpsyg.2017.00421

Bays, B. C., Visscher, K. M., Dantec, C. C. L., & Seitz, A. R. (2015). Alpha-band EEG activity in perceptual learning. Journal of Vision, 15(10), 7–7. https://doi.org/10.1167/15.10.7

Benwell, C. S. Y., Tagliabue, C. F., Veniero, D., Cecere, R., Savazzi, S., & Thut, G. (2017). Prestimulus EEG Power Predicts Conscious Awareness But Not Objective Visual Performance. ENeuro, 4(6). https://doi.org/10.1523/ENEURO.0182-17.2017

Benwell, C. S. Y., Keitel, C., Harvey, M., Gross, J., & Thut, G. (2018). Trial-by-trial covariation of pre-stimulus EEG alpha power and visuospatial bias reflects a mixture of stochastic and deterministic effects. European Journal of Neuroscience, 48(7), 2566–2584. https://doi.org/10.1111/ejn.13688

Benwell, C. S. Y., London, R. E., Tagliabue, C. F., Veniero, D., Gross, J., Keitel, C., & Thut, G. (2019). Frequency and power of human alpha oscillations drift systematically with time-on-task. Neuroimage, 192, 101–114. https://doi.org/10.1016/j.neuroimage.2019.02.067

Bernasconi, F., Manuel, A. L., Murray, M. M., & Spierer, L. (2011). Pre-stimulus beta oscillations within left posterior sylvian regions impact auditory temporal order judgment accuracy. International Journal of Psychophysiology, 79(2), 244–248. https://doi.org/10.1016/j.ijpsycho.2010.10.017

Binder, M. (2015). Neural correlates of audiovisual temporal processing – Comparison of temporal order and simultaneity judgments. Neuroscience, 300, 432–447. https://doi.org/10.1016/j.neuroscience.2015.05.011

Cecere, R., Rees, G., & Romei, V. (2015). Individual Differences in Alpha Frequency Drive Crossmodal Illusory Perception. Current Biology, 25(2), 231–235. https://doi.org/10.1016/j.cub.2014.11.034

Clayton, M. S., Yeung, N., & Kadosh, R. C. (2017). The many characters of visual alpha oscillations. European Journal of Neuroscience, 0(0). https://doi.org/10.1111/ejn.13747

Cohen, M. X. (2014). Analyzing neural time series data: Theory and practice. MIT Press.

Cohen, M. X. (2014). Fluctuations in oscillation frequency control spike timing and coordinate neural networks. Journal of Neuroscience, 34(27), 8988–8998.

Corcoran, A. W., Alday, P. M., Schlesewsky, M., & Bornkessel-Schlesewsky, I. (2018). Toward a reliable, automated method of individual alpha frequency (IAF) quantification. Psychophysiology, 55(7), e13064. https://doi.org/10.1111/psyp.13064

De Boer-Schellekens, L., Eussen, M., & Vroomen, J. (2013). Diminished sensitivity of audiovisual temporal order in autism spectrum disorder. Frontiers in Integrative Neuroscience, 7. https://doi.org/10.3389/fnint.2013.00008

Dean, C. L., Eggleston, B. A., Gibney, K. D., Aligbe, E., Blackwell, M., & Kwakye, L. D. (2017). Auditory and visual distractors disrupt multisensory temporal acuity in the crossmodal temporal order judgment task. PLOS ONE, 12(7), e0179564. https://doi.org/10.1371/journal.pone.0179564

Delorme, A., & Makeig, S. (2004). EEGLAB: An open source toolbox for analysis of single-trial EEG dynamics including independent component analysis. Journal of Neuroscience Methods, 134(1), 9–21. https://doi.org/10.1016/j.jneumeth.2003.10.009

Donohue, S. E., Green, J. J., & Woldorff, M. G. (2015). The effects of attention on the temporal integration of multisensory stimuli. Frontiers in Integrative Neuroscience, 9. https://doi.org/10.3389/fnint.2015.00032

Fister, J., Stevenson, R. A., Nidiffer, A. R., Barnett, Z. P., & Wallace, M. T. (2016). Stimulus intensity modulates multisensory temporal processing. Neuropsychologia, 88, 92–100. https://doi.org/10.1016/j.neuropsychologia.2016.02.016

Foucher, J. R., Lacambre, M., Pham, B.-T., Giersch, A., & Elliott, M. A. (2007). Low time resolution in schizophrenia: Lengthened windows of simultaneity for visual, auditory and bimodal stimuli. Schizophrenia Research, 97(1), 118–127. https://doi.org/10.1016/j.schres.2007.08.013

Gluth, S., & Meiran, N. (2019). Leave-One-Trial-Out, LOTO, a general approach to link single-trial parameters of cognitive models to neural data. ELife, 8. https://doi.org/10.7554/eLife.42607

Grabot, L., Kösem, A., Azizi, L., & Wassenhove, V. V. (2017). Prestimulus Alpha Oscillations and the Temporal Sequencing of Audiovisual Events. Journal of Cognitive Neuroscience, 29(9), 1566–1582. https://doi.org/10.1162/jocn_a_01145

Hairston, W. D., Burdette, J. H., Flowers, D. L., Wood, F. B., & Wallace, M. T. (2005). Altered temporal profile of visual–auditory multisensory interactions in dyslexia. Experimental Brain Research, 166(3), 474–480. https://doi.org/10.1007/s00221-005-2387-6

Hanslmayr, S., Aslan, A., Staudigl, T., Klimesch, W., Herrmann, C. S., & Bäuml, K.-H. (2007). Prestimulus oscillations predict visual perception performance between and within subjects. NeuroImage, 37(4), 1465–1473. https://doi.org/10.1016/j.neuroimage.2007.07.011

Iemi, L., Chaumon, M., Crouzet, S. M., & Busch, N. A. (2017). Spontaneous Neural Oscillations Bias Perception by Modulating Baseline Excitability. Journal of Neuroscience, 37(4), 807–819. https://doi.org/10.1523/JNEUROSCI.1432-16.2016

Iemi, L., & Busch, N. A. (2018). Moment-to-Moment Fluctuations in Neuronal Excitability Bias Subjective Perception Rather than Strategic Decision-Making. ENeuro, 5(3). https://doi.org/10.1523/ENEURO.0430-17.2018

Ikumi, N., Torralba, M., Ruzzoli, M., & Soto-Faraco, S. (2019). The phase of pre-stimulus brain oscillations correlates with cross-modal synchrony perception. European Journal of Neuroscience, 49(2), 150–164. https://doi.org/10.1111/ejn.14186

Keil, J., & Senkowski, D. (2017). Individual Alpha Frequency Relates to the Sound-Induced Flash Illusion. Multisensory Research, 30(6), 565–578. https://doi.org/10.1163/22134808-00002572

Klimesch, W., Doppelmayr, M., Schimke, H., & Pachinger, T. (1996). Alpha Frequency, Reaction Time, and the Speed of Processing Information. Journal of Clinical Neurophysiology, 13(6), 511–518.

Klimesch, W., Doppelmayr, M., Russegger, H., Pachinger, T., & Schwaiger, J. (1998). Induced alpha band power changes in the human EEG and attention. Neuroscience Letters, 244(2), 73–76. https://doi.org/10.1016/S0304-3940(98)00122-0

Kriegeskorte, N., Simmons, W. K., Bellgowan, P. S., & Baker, C. I. (2009). Circular analysis in systems neuroscience – the dangers of double dipping. Nature Neuroscience, 12(5), 535–540. https://doi.org/10.1038/nn.2303

Lange, J., Oostenveld, R., & Fries, P. (2013). Reduced Occipital Alpha Power Indexes Enhanced Excitability Rather than Improved Visual Perception. Journal of Neuroscience, 33(7), 3212–3220. https://doi.org/10.1523/JNEUROSCI.3755-12.2013

Lee, H., & Noppeney, U. (2011). Long-term music training tunes how the brain temporally binds signals from multiple senses. Proceedings of the National Academy of Sciences, 108(51), E1441–E1450. https://doi.org/10.1073/pnas.1115267108

Leonardelli, E., Braun, C., Weisz, N., Lithari, C., Occelli, V., & Zampini, M. (2015). Prestimulus oscillatory alpha power and connectivity patterns predispose perceptual integration of an audio and a tactile stimulus. Human Brain Mapping, 36(9), 3486–3498. https://doi.org/10.1002/hbm.22857

Lewald, J., & Guski, R. (2003). Cross-modal perceptual integration of spatially and temporally disparate auditory and visual stimuli. Cognitive Brain Research, 16(3), 468–478. https://doi.org/10.1016/S0926-6410(03)00074-0

Lou, B., Li, Y., Philiastides, M. G., & Sajda, P. (2014). Prestimulus alpha power predicts fidelity of sensory encoding in perceptual decision making. NeuroImage, 87, 242–251. https://doi.org/10.1016/j.neuroimage.2013.10.041

Love, S. A., Petrini, K., Pernet, C. R., Latinus, M., & Pollick, F. E. (2018). Overlapping but Divergent Neural Correlates Underpinning Audiovisual Synchrony and Temporal Order Judgments. Frontiers in Human Neuroscience, 12. https://doi.org/10.3389/fnhum.2018.00274

Maris, E., & Oostenveld, R. (2007). Nonparametric statistical testing of EEG- and MEG-data. Journal of Neuroscience Methods, 164(1), 177–190. https://doi.org/10.1016/j.jneumeth.2007.03.024

Martin, B., Giersch, A., Huron, C., & van Wassenhove, V. (2013). Temporal event structure and timing in schizophrenia: Preserved binding in a longer “now”. Neuropsychologia, 51(2), 358–371. https://doi.org/10.1016/j.neuropsychologia.2012.07.002

Meredith, M. A., Nemitz, J. W., & Stein, B. E. (1987). Determinants of multisensory integration in superior colliculus neurons. I. Temporal factors. Journal of Neuroscience, 7(10), 3215–3229. https://doi.org/10.1523/JNEUROSCI.07-10-03215.1987

Navarra, J., Vatakis, A., Zampini, M., Soto-Faraco, S., Humphreys, W., & Spence, C. (2005). Exposure to asynchronous audiovisual speech extends the temporal window for audiovisual integration. Cognitive Brain Research, 25(2), 499–507. https://doi.org/10.1016/j.cogbrainres.2005.07.009

Necker, L. (1832). LXI. Observations on some remarkable optical phænomena seen in Switzerland; and on an optical phænomenon which occurs on viewing a figure of a crystal or geometrical solid. The London, Edinburgh, and Dublin Philosophical Magazine and Journal of Science, 1(5), 329–337. https://doi.org/10.1080/14786443208647909

Nelli, S., Itthipuripat, S., Srinivasan, R., & Serences, J. T. (2017). Fluctuations in instantaneous frequency predict alpha amplitude during visual perception. Nature Communications, 8(1), 2071. https://doi.org/10.1038/s41467-017-02176-x

Nunez, P. L. (2000). Toward a quantitative description of large-scale neocortical dynamic function and EEG. Behavioral and Brain Sciences, 23(3), 371–398. https://doi.org/10.1017/S0140525X00003253

Oostenveld, R., Fries, P., Maris, E., & Schoffelen, J.-M. (2011). FieldTrip: Open Source Software for Advanced Analysis of MEG, EEG, and Invasive Electrophysiological Data. Intell. Neuroscience, 2011, 1:1–1:9. https://doi.org/10.1155/2011/156869

Pearson, K. (1915). On the partial correlation ratio. Proceedings of the Royal Society of London. Series A: Containing Papers of a Mathematical and Physical Character, 91(632), 492–498.

Peterson, E. J., & Voytek, B. (2018). The trade-off between neural computation and oscillatory coordination. https://doi.org/10.1101/309427

Piantoni, G., Romeijn, N., Gomez-Herrero, G., Werf, Y. D. V. D., & Someren, E. J. W. V. (2017). Alpha Power Predicts Persistence of Bistable Perception. Scientific Reports, 7(1), 5208. https://doi.org/10.1038/s41598-017-05610-8

Powers, A. R., Hillock, A. R., & Wallace, M. T. (2009). Perceptual Training Narrows the Temporal Window of Multisensory Binding. Journal of Neuroscience, 29(39), 12265–12274. https://doi.org/10.1523/JNEUROSCI.3501-09.2009

Prins, N., & Kingdom, F. A. A. (2018). Applying the Model-Comparison Approach to Test Specific Research Hypotheses in Psychophysical Research Using the Palamedes Toolbox. Frontiers in Psychology, 9. https://doi.org/10.3389/fpsyg.2018.01250

Quenouille, M. H. (1949). Approximate tests of correlation in time-series 3. Mathematical Proceedings of the Cambridge Philosophical Society, 45(3), 483–484. https://doi.org/10.1017/S0305004100025123

Richter, C. G., Thompson, W. H., Bosman, C. A., & Fries, P. (2015). A jackknife approach to quantifying single-trial correlation between covariance-based metrics undefined on a single-trial basis. NeuroImage, 114, 57–70. https://doi.org/10.1016/j.neuroimage.2015.04.040

Romei, V., Brodbeck, V., Michel, C., Amedi, A., Pascual-Leone, A., & Thut, G. (2008). Spontaneous Fluctuations in Posterior α-Band EEG Activity Reflect Variability in Excitability of Human Visual Areas. Cerebral Cortex, 18(9), 2010–2018. https://doi.org/10.1093/cercor/bhm229

Ronconi, L., Busch, N. A., & Melcher, D. (2018). Alpha-band sensory entrainment alters the duration of temporal windows in visual perception. Scientific Reports, 8(1). https://doi.org/10.1038/s41598-018-29671-5

Samaha, J., & Postle, B. R. (2015). The Speed of Alpha-Band Oscillations Predicts the Temporal Resolution of Visual Perception. Current Biology, 25(22), 2985–2990. https://doi.org/10.1016/j.cub.2015.10.007

Sauseng, P., Klimesch, W., Gerloff, C., & Hummel, F. C. (2009). Spontaneous locally restricted EEG alpha activity determines cortical excitability in the motor cortex. Neuropsychologia, 47(1), 284–288. https://doi.org/10.1016/j.neuropsychologia.2008.07.021

Savitzky, A., & Golay, M. (1964). A. Savitzky and MJE Golay, Anal. Chem. 36, 1627 (1964). Anal. Chem., 36, 1627.

Schneider, W., Eschman, A., & Zuccolotto, A. (2002). E-Prime reference guide. Psychology Software Tools, Incorporated.

Senkowski, D., Talsma, D., Grigutsch, M., Herrmann, C. S., & Woldorff, M. G. (2007). Good times for multisensory integration: Effects of the precision of temporal synchrony as revealed by gamma-band oscillations. Neuropsychologia, 45(3), 561–571. https://doi.org/10.1016/j.neuropsychologia.2006.01.013

Shen, L., Han, B., Chen, L., & Chen, Q. (2018). Perceptual inference employs intrinsic alpha frequency to resolve perceptual ambiguity. https://doi.org/10.1101/399840

Stahl, J., & Gibbons, H. (2004). The application of jackknife-based onset detection of lateralized readiness potential in correlative approaches. Psychophysiology, 41(6), 845–860. https://doi.org/10.1111/j.1469-8986.2004.00243.x

Stevenson, R. A., Zemtsov, R. K., & Wallace, M. T. (2012). Individual differences in the multisensory temporal binding window predict susceptibility to audiovisual illusions. Journal of Experimental Psychology: Human Perception and Performance, 38(6), 1517–1529. http://dx.doi.org/10.1037/a0027339

Stevenson, R. A., & Wallace, M. T. (2013). Multisensory temporal integration: Task and stimulus dependencies. Experimental Brain Research. Experimentelle Hirnforschung. Experimentation Cerebrale, 227(2), 249–261. https://doi.org/10.1007/s00221-013-3507-3

Stevenson, R. A., Siemann, J. K., Schneider, B. C., Eberly, H. E., Woynaroski, T. G., Camarata, S. M., & Wallace, M. T. (2014). Multisensory Temporal Integration in Autism Spectrum Disorders. The Journal of Neuroscience, 34(3), 691–697. https://doi.org/10.1523/JNEUROSCI.3615-13.2014

Stevenson, R. A., Park, S., Cochran, C., McIntosh, L. G., Noel, J.-P., Barense, M. D., Ferber, S., & Wallace, M. T. (2017). The associations between multisensory temporal processing and symptoms of schizophrenia. Schizophrenia Research, 179, 97–103. https://doi.org/10.1016/j.schres.2016.09.035

Strüber, D., & Herrmann, C. S. (2002). MEG alpha activity decrease reflects destabilization of multistable percepts. Cognitive Brain Research, 14(3), 370–382. https://doi.org/10.1016/S0926-6410(02)00139-8

Talsma, D., Senkowski, D., & Woldorff, M. G. (2009). Intermodal attention affects the processing of the temporal alignment of audiovisual stimuli. Experimental Brain Research, 198(2), 313–328. https://doi.org/10.1007/s00221-009-1858-6

Tukey, J. (1958). Bias and confidence in not quite large samples. Ann. Math. Statist., 29, 614.

van Driel, J., Knapen, T., van Es, D. M., & Cohen, M. X. (2014). Interregional alpha-band synchrony supports temporal cross-modal integration. NeuroImage, 101, 404–415. https://doi.org/10.1016/j.neuroimage.2014.07.022

van Eijk, R. L. J., Kohlrausch, A., Juola, J. F., & van de Par, S. (2008). Audiovisual synchrony and temporal order judgments: Effects of experimental method and stimulus type. Perception & Psychophysics, 70(6), 955–968. https://doi.org/10.3758/PP.70.6.955

van Viegen, T., Charest, I., Jensen, O., & Mazaheri, A. (2017). Alpha/beta power compresses time in sub-second temporal judgments. https://doi.org/10.1101/224386

VanRullen, R. (2016). Perceptual Cycles. Trends in Cognitive Sciences, 20(10), 723–735. https://doi.org/10.1016/j.tics.2016.07.006

Von Stein, A., Rappelsberger, P., Sarnthein, J., & Petsche, H. (1999). Synchronization between temporal and parietal cortex during multimodal object processing in man. Cerebral Cortex, 9(2), 137–150. Scopus.

von Stein, A., & Sarnthein, J. (2000). Different frequencies for different scales of cortical integration: From local gamma to long range alpha/theta synchronization. International Journal of Psychophysiology, 38(3), 301–313. https://doi.org/10.1016/S0167-8760(00)00172-0

Wallace, M. T., & Stevenson, R. A. (2014). The construct of the multisensory temporal binding window and its dysregulation in developmental disabilities. Neuropsychologia, 64, 105–123. https://doi.org/10.1016/j.neuropsychologia.2014.08.005

Watkins, S., Shams, L., Tanaka, S., Haynes, J.-D., & Rees, G. (2006). Sound alters activity in human V1 in association with illusory visual perception. NeuroImage, 31(3), 1247–1256. https://doi.org/10.1016/j.neuroimage.2006.01.016

Wutz, A., Weisz, N., Braun, C., & Melcher, D. (2014). Temporal Windows in Visual Processing: ‘Prestimulus Brain State’ and ‘Poststimulus Phase Reset’ Segregate Visual Transients on Different Temporal Scales. Journal of Neuroscience, 34(4), 1554–1565. https://doi.org/10.1523/JNEUROSCI.3187-13.2014

Wutz, Andreas, Melcher, D., & Samaha, J. (2018). Frequency modulation of neural oscillations according to visual task demands. Proceedings of the National Academy of Sciences, 115(6), 1346–1351. https://doi.org/10.1073/pnas.1713318115

Yuan, X., Li, H., Liu, P., Yuan, H., & Huang, X. (2016). Pre-stimulus beta and gamma oscillatory power predicts perceived audiovisual simultaneity. International Journal of Psychophysiology, 107, 29–36. https://doi.org/10.1016/j.ijpsycho.2016.06.017

